# Liquid-phase determination of Arabidopsis respiration and photosynthesis using Clark-type O_2_ electrodes

**DOI:** 10.1101/2025.05.04.652138

**Authors:** Florencia Sena, Camila Couture, Andrés Berais-Rubio, A. Harvey Millar, Santiago Signorelli

## Abstract

Photosynthesis and respiration are fundamental metabolic processes in plants, tightly connected through shared substrates, energy dynamics, and redox balance. Arabidopsis is the key genetic model for plants but monitoring these sorts of physiological processes presents significant challenges using traditional gas-exchange or fluorescence-based techniques due to the small size of intact *Arabidopsis thaliana* (arabidopsis) seedlings. Here, we validate and characterize the use of Clark-type oxygen electrodes, specifically the Hansatech Oxytherm+P system, to quantify both photosynthetic and respiratory activity in intact arabidopsis seedlings. By monitoring oxygen evolution in dark and light phases, we demonstrate that oxygen consumption and production correspond to mitochondrial respiration and photosynthesis, respectively. These processes were modulated by tissue biomass, light intensity, developmental stage, and stress conditions. Specific inhibitors such as potassium cyanide and paraquat confirmed that the recorded changes in oxygen concentrations reflected mitochondrial cytochrome oxidase activity and photosystem electron transport-dependent oxygen production, respectively. Moreover, oxygen evolution increased significantly with bicarbonate supplementation, validating the system’s sensitivity to carbon fixation. We further showed that photosynthetic activity measured with this method correlates with a quantitative green index and responds dynamically to de-etiolation, abiotic stress (salt, osmotic, oxidative), and temperature shifts. Our study lays the groundwork for measuring photosynthesis based on oxygen evolution and respiration in arabidopsis knockout mutants, CRISPR lines, overexpression lines and ecotypes using Clark-type oxygen electrodes and highlights key considerations and limitations to consider when applying this approach. This platform could also be adapted for many other small tissue plant samples.

## 1. INTRODUCTION

### 1.1. Measuring photosynthesis and respiration in small plant systems

Photosynthesis and respiration are two fundamental and tightly interconnected metabolic processes in plants. While photosynthesis converts light energy into chemical energy, producing sugars and oxygen, mitochondrial respiration breaks down these organic molecules to generate ATP, consuming oxygen and releasing carbon dioxide. Together, these pathways determine carbon balance, biomass accumulation, and overall plant performance, and their measurement is essential in plant physiology, stress biology, and crop improvement research (Amthor et al., 2019).

However, quantifying these processes in small or early-stage plant tissues, such as arabidopsis seedlings, presents methodological challenges. Conventional tools like infrared gas analyzers (IRGA) and chlorophyll fluorescence-based devices (e.g., PAM fluorometers) often require a minimum tissue size, are not easily adaptable to roots or etiolated tissues, and may not provide direct rates of oxygen production or consumption (Walker et al., 2024; Calzadilla et al., 2022). Arabidopsis research often involves growing plant seedlings on agar plates or in liquid media and assessing whole plant phenotypes (Rivero et al., 2014; Signorelli et al., 2016). This creates a gap in accessible methods for determining respiration and photosynthesis in small model systems, which are widely used in genetic and developmental studies.

### 1.2. Oxygen-based approaches and the use of Clark-type electrodes to assess plant photosynthesis

Oxygen exchange (production during photosynthesis and consumption during respiration) offers a direct proxy for metabolic activity and has been widely used on photosynthetic organisms in the last decades (Hunt, 2003; Takagi et al., 2016). The method was initially developed in the 1950s by Leland Clark for monitoring blood O_2_ tensions during cardiac surgery (Clark et al., 1953) via its electrochemical reduction at a platinum cathode. Historically, the possibility of evaluating oxygen consumption by Clark-type electrodes to determine cellular respiration has been widely reported in the literature on different organisms such as animals, bacteria, and plants (Silva and Oliveira., 2012, Shirato et al., 2024, Oh et al., 2022). While cellular respiration has been widely studied using these types of electrodes (Lee et al., 2024, Oh et al., 2023, Oh et al., 2022), literature reports of photosynthesis measurements were mainly limited to cyanobacteria (Ungerer et al., 2018), unicellular algae (Ananyev et al., 2016, Kosourov et al., 2018, Vera-Vives et al., 2024), mesophyll protoplasts (Hebbelmann et al., 2012) or eventually isolated thylakoid membranes (Fitzpatrick et al., 2022) from higher plants. Clark-type O_2_ electrodes are very rarely reported to determine photosynthesis in intact higher plants.

Recent improvements in commercial Clark-type electrode systems, such as the Oxytherm+P (Hansatech), now allow simultaneous control of temperature and illumination, opening the possibility of assessing both photosynthesis and respiration in whole plant seedlings. In a recent study, some of us used a Clark-type electrode systems complemented with an external source of light to measure photosynthetic activity in 7-day-old arabidopsis seedlings (Wijerathna-Yapa et al., 2021), a material not easily suited for IRGA-based gas exchange. This suggests that Clark-type electrodes could offer a valuable, cost-effective, and flexible approach for studying changes in plant metabolism in small tissues.

In this study, we evaluate the use of the new Oxytherm+P Clark-type oxygen electrode to measure both respiration and photosynthesis in intact arabidopsis seedlings. We assess the system’s responsiveness to biomass quantity, light intensity, developmental stage, and abiotic stresses, and validate its specificity using pathway inhibitors and bicarbonate supplementation. Furthermore, we identify practical considerations and limitations for its broader application in plant physiology studies.

## 2. MATERIAL AND METHODS

### 2.1. Plant material and growth conditions

*A. thaliana* wild-type plants from ecotype Columbia-0 (Col-0) were obtained from the Arabidopsis Biological Resource Center and are referred to generically as arabidopsis. For plant growth assays on plates, arabidopsis seeds were surface sterilized with 70 % EtOH for 2 min, followed by 10 % NaClO and 0.5 % Tween 20 for three min, then washed three times for three min in sterile deionized water. Seeds were placed on 0.5 x MS (Murashige and Skoog., 1962) plates containing 0.5 % sucrose and 1.2 % plant agar, and stratified in darkness for two days at 4 °C, followed by pre-germination under long-day conditions (16 h/ 8 h light/dark, 22 °C, light intensity ≈ 100 μmol m^-2^ s^-1^, and relative humidity 40 %) for seven days.

### 2.2. General protocol for oxygen evolution measurement on *A. thaliana* seedlings

Oxygen evolution was measured using highly sensitive S1 Clark-type polarographic oxygen electrodes (Hansatech Oxytherm+P chamber). The oxygen electrodes were assembled with the electrodes immersed in 50 % saturated potassium chloride solution beneath an oxygen-permeable PTFE membrane. A paper spacer was placed underneath the membrane to provide a uniform electrolyte layer between the anode and cathode. Two oxygen electrodes were used simultaneously to measure oxygen evolution in different samples. The electrodes were calibrated before measurements using air-saturated and deoxygenated (dithionite) water to set 100 %- and 0 %-oxygen levels. The electronic electrode chamber with solid-state Peltier was set to 25 °C and stirrer speed at 70 rpm, and the liquid phase consisted of a buffer (1 mM NaHCO_3_, 10 mM HEPES-KOH, pH 7.5), freshly prepared the same day of each measurement.

Seven-day-old seedlings were transferred to the electrode chamber containing two ml of the buffer HEPES with NaHCO_3_. Seedlings were gently submerged but not allowed to touch the electrode membrane. Before each measurement, seedlings were blotted dry and scale weighted.

During the measurements we defined three different phases. The first phase consisted of three to five min of measurement under dark conditions to determine oxygen consumption (cellular respiration). A sliding shutter plate on the chamber allows the sample to be placed in full dark conditions for dark respiration measurement phases. After at least three minutes of linear slope, the second phase was initiated by turning on the light, and oxygen evolution was recorded for at least five additional minutes. Then, the light was turned off again, and another three to five minutes were recorded in darkness (third phase).

Oxygen consumption and production rates were determined as:

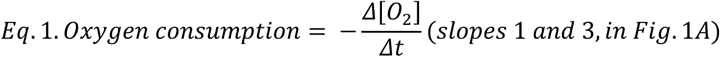

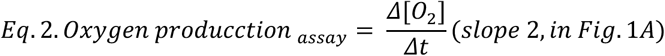

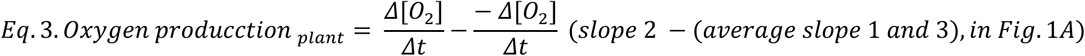

Respiration was calculated as the average of the oxygen consumption determined from the slopes of phases one and three (light off) (Eq. 1) per FW of sample and expressed as nmol O_2_ min^-1^ g^-1^ FW. Light-dependent oxygen production (i.e., photosynthesis) was calculated as the difference between the slope obtained in light condition (phase two) and darkness (phases one and three) (Eq. 3) and expressed as nmol O_2_ min^-1^ g^-1^ FW.

### 2.3. Respiration and photosynthesis inhibition assays

Seven-day-old arabidopsis seedlings were used for oxygen evolution measurement as described in section 2.2 with the following modifications. For the respiration inhibition assay, we determined two dark phases, the first one as described in 2.2, while the second one was initiated by the addition of potassium cyanide (KCN) into the buffer to a final concentration of 3 mM. For the photosynthesis inhibition assay, was developed within three phases, the first two phases as described in section 2.2, but the last phase was in light conditions with the addition of paraquat to a final concentration of 100 μM.

### 2.4. Bicarbonate treatment

Seven-day-old arabidopsis seedlings were used for oxygen evolution measurement as described in section 2.2 with the following modifications. During the first and second phases of measuring, the seedlings were placed into the chamber without the addition of bicarbonate (NaHCO_3_). However, in the third phase, under light conditions, 20 μL of a freshly prepared stock solution of 100 mM of NaHCO_3_ were added. A fourth phase of dark condition was included to determine respiration (representing the third phase in the method described 2.2).

### 2.5. Light intensity treatments

A week-old arabidopsis seedlings were exposed to different light intensities to determine oxygen production over time. The Oxytherm+P electrode chamber is fitted with two high-intensity automated white LED light sources mounted against the outer wall of the reaction vessel to provide a uniform illumination of the seedlings. We have used the 20 individual light steps configured within the OxyTrace+ software to calibrate the LED photon flux density (PFD). The light meter and thermometer QTP1 PAR/temperature sensor was connected to the rear of the Oxytherm+P control unit and placed into the reaction vessel before the addition of any liquids for the quantification.

### 2.6. Temperature treatments

For determining the oxygen evolution at different temperatures, the oxygen electrode was calibrated at five different temperatures (5, 15, 25, 35, and 40 °C). The calibrations were performed as described in section 2.2. After calibration at one temperature, the seedlings were placed into the chamber to proceed with the oxygen evolution measurement as described in section 2.2.

### 2.7. De-etiolation and greening treatments

The developmental transition of dark-to-light greening of arabidopsis seedlings was used to detect changes in oxygen evolution. Photoautotrophic seedlings were obtained from arabidopsis seeds grown seven days in the dark condition and then grown under a 16/8 h light/dark period with a light intensity of ≈ 100 μmol m^−2^ sec^−1^ at 22 °C for 0, 12, 24, 48, or 168 hours further. Plates were photographed each time, and the green index (GI) was calculated before the oxygen measurement.

### 2.8. Abiotic stress treatments

Different abiotic stresses (osmotic, salinity, and oxidative) were applied to arabidopsis seedlings to evaluate the oxygen evolution. Sterile arabidopsis seedlings were pre-germinated on 0.5 x MS medium for five days and then transferred to the different stressed medium for two days. Stress treatments were induced by adding 150 mM NaCl, 200 mM mannitol, or 1 M hydrogen peroxide to the 0.5 x MS medium plates. The oxygen measurements were recorded on 2 ml of buffer HEPES and NaHCO_3_ with the fresh addition of 150 mM NaCl, 200 mM mannitol, or 1 M hydrogen peroxide.

### 2.9. Green index (GI) determination

To assess the greening of seedlings during the light-to-dark transition or under different stress conditions and its correlation with photosynthetic capacity, we calculated the Green Index (GI) as described in Signorelli et al., 2023. Briefly, a camera phone was used to obtain the pictures of seedlings. Leaf green areas were evaluated using the Raw Therapee open-source software to get the RGB values. GI was calculated from the RGB values using the Green Index software is available at www.foodandplantbiology.fagro.edu.uy or www.foodandplantbiology.com.

### 2.10. Statistical analysis

Analysis of variance were performed with data from at least three replicates in all the cases and the means were compared using Tukey’s post-hoc honest significant difference test at the P < 0.05 level using the open software R. When statistical comparisons between two groups was needed, the student’s t-test were performed using R. Differences were considered statistically significant at *p* < 0.05. For some continuous variables we represented the adjusted regression for the independent measurements and a grey area representing the 95 % confidence level intervals for the regression prediction using the loess method in the ggplot2 package of R.

## 3. RESULTS

### 3.1. Validation of respiration and photosynthesis measurements via oxygen evolution in arabidopsis seedlings

We used a liquid-phase Clark-type oxygen electrode to monitor the oxygen evolution in arabidopsis seedlings under alternating dark and light conditions. Oxygen consumption during the dark phase (slopes 1 and 3; **Figure 1A**) was used to calculate the rate of mitochondrial respiration. We estimated photosynthetic activity by subtracting the average darkness-phase oxygen consumption from the light-phase oxygen production (slope 2), providing a measure of net photosynthesis (Equation 3; **Figure 1A**).

To assess whether respiration and photosynthesis rates scale linearly with sample size, we measured oxygen evolution in seedling clumps ranging from 20 to 200 mg fresh weight. Both respiration and photosynthesis showed a strong linear correlation with fresh weight (**Figure 1B** and **Figure 1C**). This confirmed that the system reliably detects proportional metabolic activity across this biomass range and allowed us to normalize all subsequent measurements to fresh weight.

**Figure 1.**
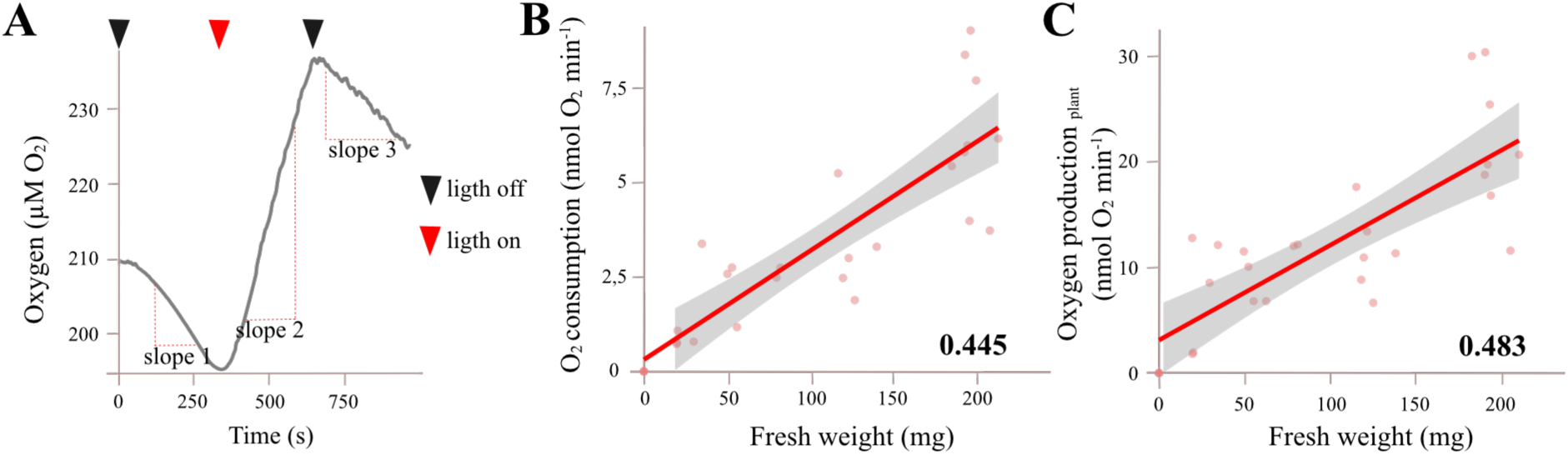
Determination of respiration and photosynthesis on intact arabidopsis plants by monitoring O2 evolution with a Clark-type electrode. **A.** Representation of oxygen evolution during respiration and photosynthesis measurements. Slopes 1 and 3 were determined during the absence of light (dark triangle) and used to calculate respiration rates. Slope 2 was determined during the presence (red triangle) of light and used together with respiration rates to determine plant oxygen production (Eq. 3, i.e. oxygen production rate is equal to Slope 2 minus oxygen consumption rate). **B.** Relationship between O2 consumption (Eq. 1), as an indicator of plant respiration, and plant fresh weight. The shaded area shows the 95 % confidence interval and the Pearson correlation coefficient for these parameters is given in bold numbers. **C.** Relationship between oxygen production (Eq. 3), as an indicator of plant photosynthesis, and plant fresh weight. The shaded area shows the 95 % confidence interval and the Pearson correlation coefficient for these parameters is given in bold numbers.

To verify that the recorded oxygen signals corresponded specifically to mitochondrial respiration and chloroplast photosynthesis, we used pathway-specific inhibitors for each metabolic process. The addition of 3 mM potassium cyanide (KCN), a cytochrome C oxidase inhibitor, during the dark phase reduced oxygen consumption by approximately 60 % (**Figure 2A**), indicating that most of the oxygen consumption is due to cytochrome C oxidase-dependent respiration. The residual oxygen consumption may reflect cyanide-insensitive alternative oxidase activity or other oxygen-consuming processes.

Similarly, to validate the light-phase oxygen evolution as photosynthetic, we treated seedlings with 100 µM paraquat (1,1-dimethyl-4,4′-bipyridinium dichloride), an herbicide that inhibits photosynthesis by accepting electrons from photosystem I (PSI) at the thylakoid membrane of the chloroplast, blocking the electron transport chain. This treatment resulted in a significant (≈ 75%) decrease in oxygen production during the light phase (**Figure 2B**), confirming that the measured oxygen evolution is primarily due to photosynthetic activity.

**Figure 2.**
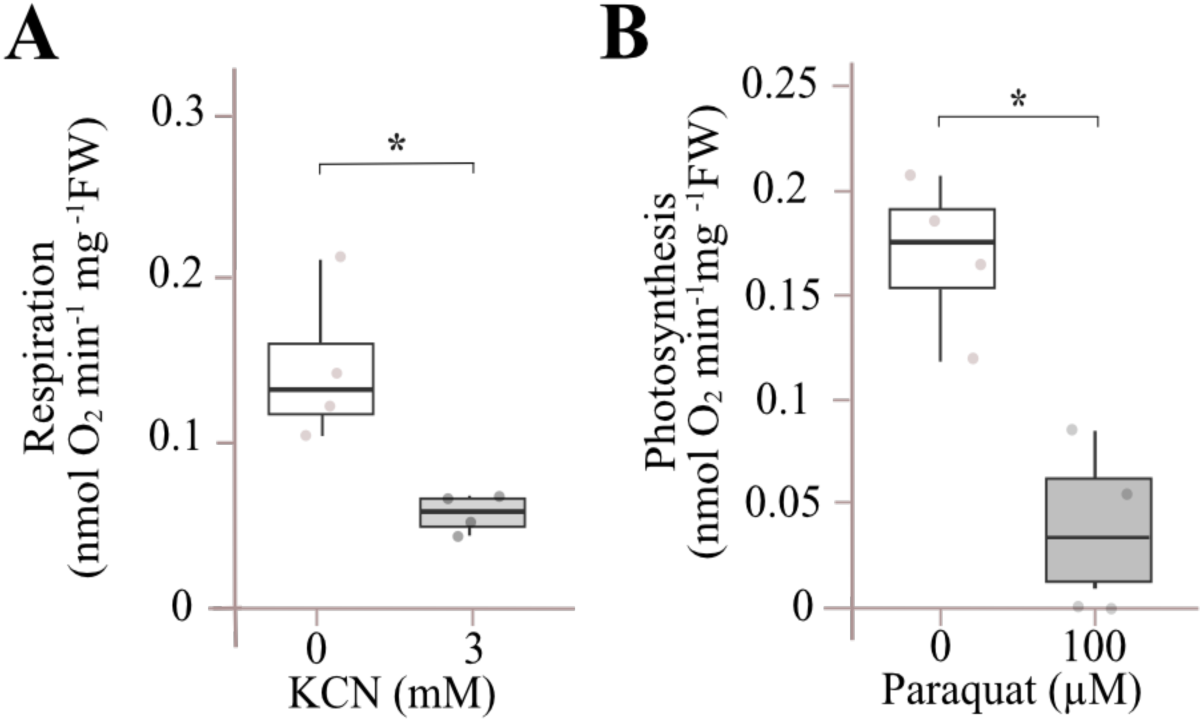
Oxygen production and consumption by arabidopsis seedlings in the presence of inhibitors for mitochondrial respiratory chain and photosynthetic electron transport. **A.** Respiration in seven-old arabidopsis seedlings with and without 3 mM KCN (cytochrome oxidase-dependent respiration inhibitor). **B.** Photosynthesis in seven days-old arabidopsis seedlings with and without 100 µM Paraquat (photosynthetic electron transfer inhibitor). Boxes represent the mean, horizontal lines the median and vertical lines the standard deviation. Dots represent each independent replicate. Asterisks represent significant differences according to t-student (p < 0.05, n = 4).

### 3.2. Bicarbonate effect on photosynthetic activity

Our previous results confirmed that light-dependent oxygen production is largely inhibited by Paraquat, indicating it originates from photosynthetic activity. However, since water splitting can occur in the absence of carbon fixation (Renger and Renger, 2008), we further tested whether the oxygen evolution measured under light is influenced by carbon availability. A known limitation of photosynthesis measurements in liquid phase is the reduced availability of CO_2_ due to its low solubility and diffusion rate in water, which can constrain carbon fixation and consequently oxygen evolution (Baker, 2008). Therefore, we compared oxygen production in seedlings with and without supplementation of 1 mM sodium bicarbonate (NaHCO_3_), an inorganic carbon source. This approach allows not only to assess the extent of carbon limitation but also provides a practical means to enhance photosynthetic rates in liquid-phase measurements if required.

The addition of bicarbonate significantly enhanced the rate of oxygen evolution under light conditions (**Figure 3A**), resulting in approximately a six-fold increase in photosynthetic activity compared to control (**Figure 3C**). A positive oxygen evolution rate was still observed in the absence of added bicarbonate, but because the buffer was not CO_2_-depleted and atmospheric CO_2_ concentrations in water at room temperature are about 13.7 µM (Wiebe and Gaddy, 1940), we cannot rule out carbon fixation entirely in this condition.

These results indicate that oxygen production measured using the Oxytherm+P system is not only light-dependent, but also responsive to carbon fixation capacity. This reinforces the potential of this method to serve as a proxy for photosynthetic activity in intact seedlings.

**Figure 3.**
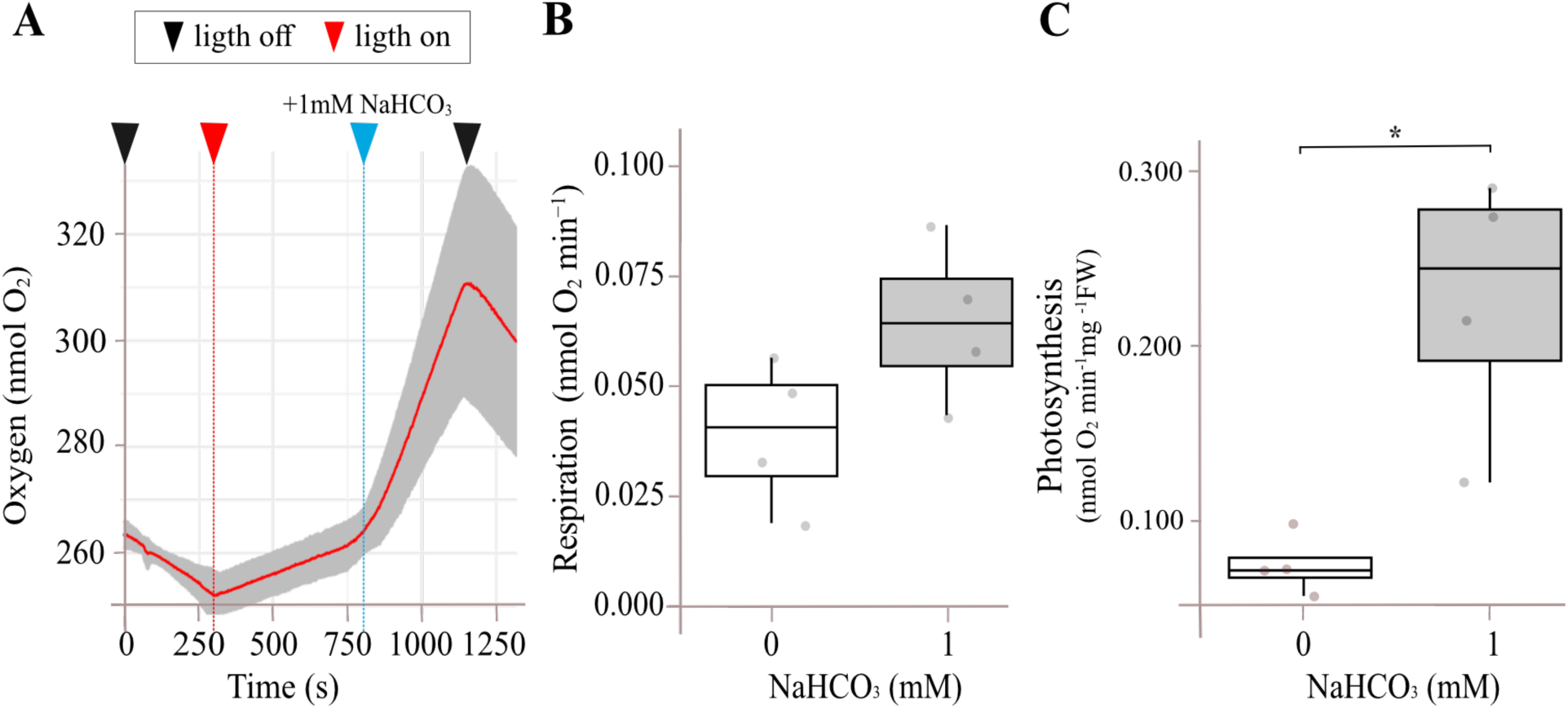
Dependence of oxygen consumption and production by arabidopsis seedlings with carbon dioxide supply. **A.** Effect of the addition of NaCO_3_ on O2 evolution over time. **B.** Respiration and **C.** Photosynthesis in seven days-old arabidopsis seedlings without and with 1 mM NaCO_3_. Boxes represent the mean, horizontal lines the median and vertical lines the standard deviation. Dots represent each independent replicate. Asterisk represents significant differences according to t-student (p <0.05, n = 4).

### 3.3. Effect of light intensity on photosynthetic oxygen evolution

To characterize the light response of photosynthesis under our experimental setup, we first calibrated the light output of the Oxytherm+P system. While the manufacturer reports that the LED light source can reach up to 4000 µmol photons m^-2^ s^-1^, our measurements using a QTP1 PAR/temperature sensor placed inside the chamber showed that the system can deliver up to 4500 µmol photons m^-2^ s^-1^ at 100% output. Light intensity increased linearly with software-controlled percentage settings (**Figure 4A**).

We then examined the effect of light intensities on photosynthetic activity, measured as oxygen evolution. The light response curve of Arabidopsis seedlings showed a linear increase in oxygen production up to approximately 400 µmol photons m^-2^ s^-1^, followed by saturation at around 600µmol photons m^-2^ s^-1^ (**Figure 4B**. This response is consistent with previously reported light saturation thresholds for arabidopsis photosynthesis (Popova et al., 2018).

**Figure 4.**
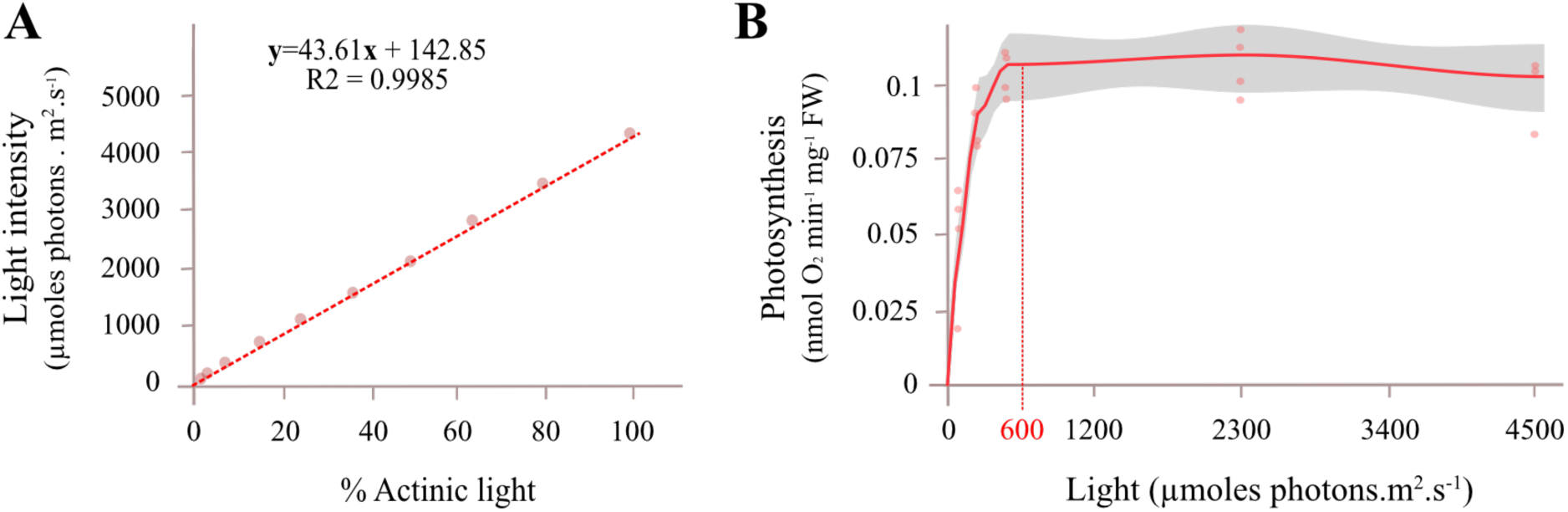
Relationship between photosynthetic activity determined by oxygen evolution and the intensity of the light source. **A.** Determination of the intensity of the Oxytherm+P electrode light source at different percentages. The QTP1 PAR/temperature sensor connected to the Oxytherm+P control unit and placed into the reaction vessel allowed us to determine the corresponding Photon Flux Density (PFD) from the % of actinic light of the white LED light source. **B.** Photosynthesis of arabidopsis seedlings in response to different light intensities. Seedlings were exposed to different light intensities (from 0 to 4500 µmol photons m^-2^ s^-1^) and photosynthesis was recorded. The light red dots represent the independent replicates (n =4), the red line represents adjusted regression for the independent measurements and the grey area represents the 95 % confidence level intervals for the regression prediction.

### 3.4. Effect of temperature on respiration and photosynthesis

Temperature influences the rate of both chemical and enzymatic reactions and thus modulates key physiological processes such as respiration and photosynthesis. To evaluate the effect of temperature under our experimental conditions, we measured oxygen evolution in arabidopsis seedlings at temperatures ranging from 5 to 40 °C using the Oxytherm+P system.

As expected, both respiration and photosynthesis rates were significantly reduced at low temperatures compared to the control condition (25 °C, **Figure 5**). Oxygen evolution increased progressively with temperature, reaching maximum values at 35 °C and no statistically significant difference was observed between 35 °C and 40 °C (**Figure 5**), which was the upper limit of the chamber’s temperature control. These results confirm that the system can capture temperature-dependent changes in metabolic activity and highlight the thermal sensitivity of both photosynthetic and respiratory processes in young arabidopsis seedlings.

**Figure 5.**
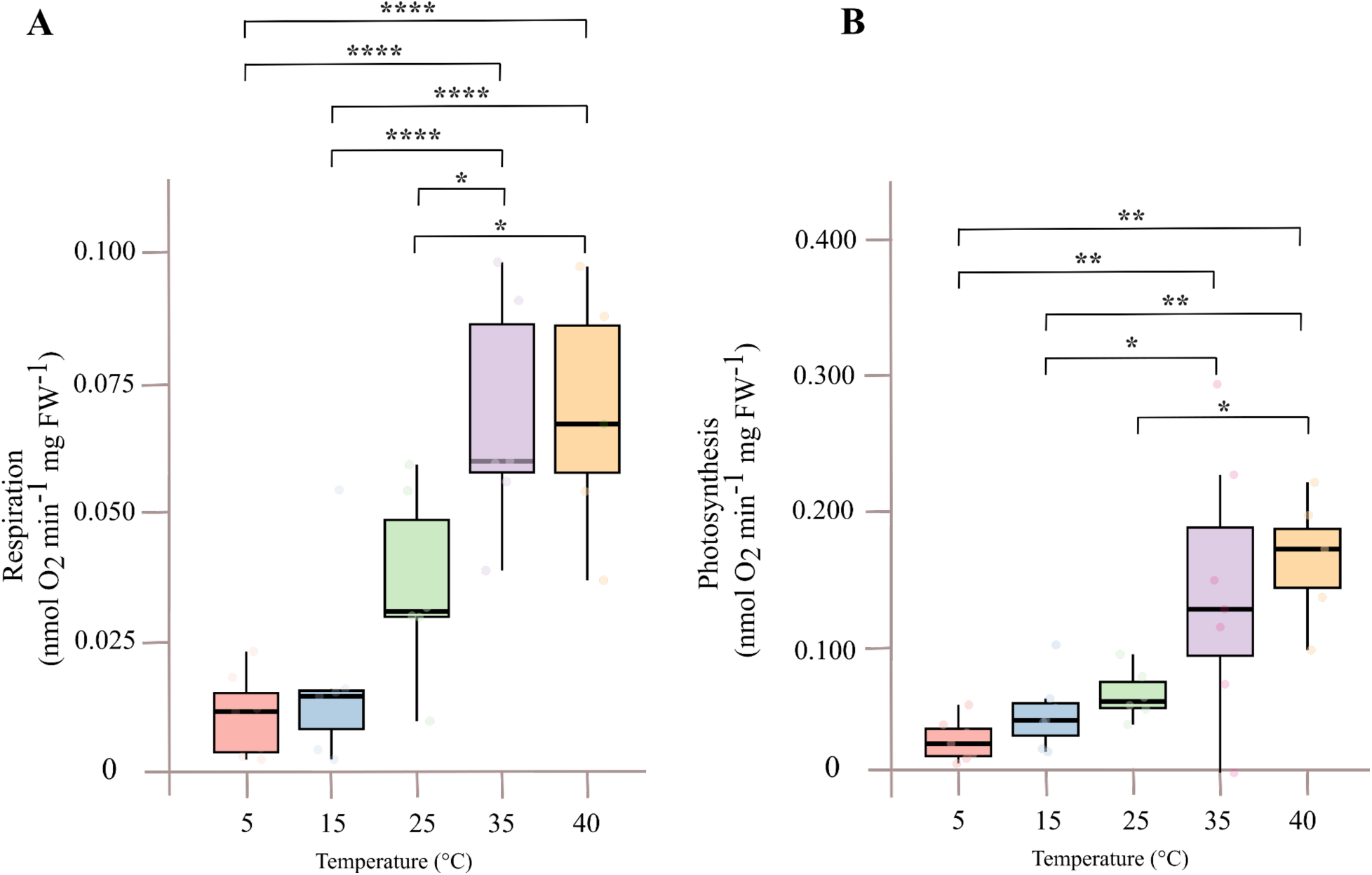
Impact of temperature on oxygen consumption and evolution rate from arabidopsis seedlings. **A.** Respiration. **B.** Photosynthesis. Boxes represent the interquartile range (IQR), which spans from the first quartile (Q1) to the third quartile (Q3). The horizontal line inside the box indicates the median. Whiskers extend to the furthest data points within 1.5 times the IQR from the quartiles. Data points beyond this range are shown as individual dots and represent outliers. Dots represent the independent replicates. Asterisks represent significant differences according to ANOVA test (* p < 0.05, ** p < 0.01, and *** p < 0.001, n ≈ 4).

### 3.5. Photosynthesis determined by oxygen evolution in etiolated, de-etiolated and non-etiolated seedlings

We used etiolation and de-etiolation processes (greening) to test the capacity of photosynthetic measurements by oxygen evolution to identify differences in photosynthetic capacity during seedling development. Arabidopsis seedlings were grown in complete darkness for seven days (0 h, etiolated), or transferred to light for 6, 24 and 48 h (de-etiolated). As a control, non-etiolated seedlings were grown under standard photoperiod conditions for seven days (168 h) (**Figure 6A**). As expected, etiolated seedlings showed elongated hypocotyls, closed apical hook and pallid yellow cotyledons. Upon light exposure, cotyledons gradually opened and became visibly greener (Figure 6B).

To quantify this greening, we quantified the Green Index, a parameter based on RGB image analysis shown to correlate with chlorophyll content (Signorelli et al. 2023). GI values increased progressively during de-etiolation, reaching levels comparable to fully green seedlings after 24h of light exposure (**Figure 6C**), confirming the effectiveness of our light treatments in inducing chloroplast development.

We next measured photosynthetic activity by oxygen evolution. As expected, etiolated seedlings displayed no detectable photosynthetic activity, and only a slight, not statistically significant, increase was observed at 6 and 24 h of de-etiolation treatment. By 48 h, photosynthetic rates increased markedly, though they remained lower than those in fully green seedlings grown under standard conditions (**Figure 6E**). It is worth mentioning that, in our calculation, photosynthesis is expressed relative to the total fresh weight of seedlings. In de-etiolated seedlings the portion of photosynthetic tissue is smaller than in seedlings grown under control conditions, meaning that we cannot conclude that cotyledons at 48 h are less active photosynthetically than those at 168 h.

Finally, we found a strong correlation between GI and photosynthetic activity across treatments (**Figure 6F**), supporting the use of GI as a simple proxy for photosynthetic potential. These results confirmed that our method can detect functional differences in photosynthesis during early photomorphogenesis in arabidopsis.

**Figure 6.**
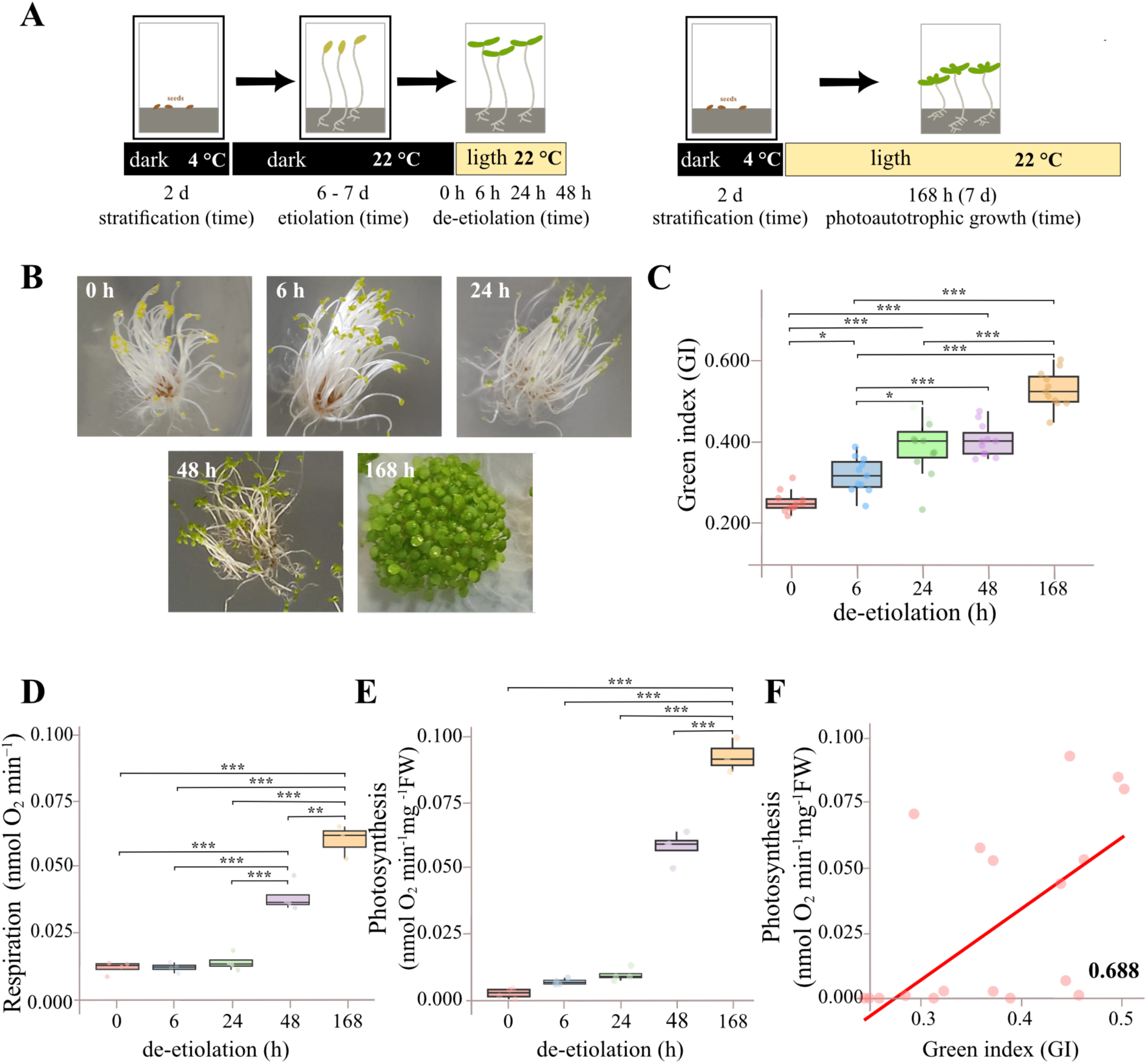
Effect of different light treatments on photosynthetic and respiratory capacity determined by oxygen evolution and consumption. **A.** Experimental design to obtain seedlings with different greening. Seeds were germinated and placed on MS medium in darkness for two days at 4 °C. For etiolation, seeds were placed at 21 °C and kept in darkness for six-seven days. For de-etiolation, etiolated seedlings were exposed to light for 6 h, 24 h, and 48 h. For non-etiolated seedlings, seeds were placed at 21°C under light condition with a 16/8 light/dark photoperiod during 168 h. **B.** Phenotypes of arabidopsis seedlings after 0, 6, 24, 48, and 168 h of light exposure. **C.** Green index of seedling exposed to different light treatments. Asterisks represent significant differences according to ANOVA test (* p < 0.05, ** p < 0.01, and *** p < 0.001, n ≈ 10). **D.** Respiration and **E.** Photosynthetic capacity of seedlings at different stages of the photomorphogenic process determined by oxygen evolution. Asterisks represent significant differences according to ANOVA test (* p < 0.05, ** p < 0.01, and *** p < 0.001, n ≈ 3). **F.** Correlation between GI and photosynthesis. Pearson correlation coefficient is presented in bold in the right bottom corner.

### 3.5. Impact of abiotic stress on photosynthesis and respiration

Abiotic stresses such as salinity, osmotic imbalance, and oxidative damage are known to affect plant metabolism, including both photosynthesis and respiration (Flexas et al., 2005, Muhammad et al., 2021). To test the sensitivity of the oxygen evolution-based method for assessing photosynthesis under abiotic stress conditions, we exposed arabidopsis seedlings to 200 mM mannitol (osmotic stress), 150 mM NaCl (salt stress), and 1 M hydrogen peroxide (oxidative stress) for two days prior to measurement.

Hydrogen peroxide-treated seedlings displayed clear signs of damage, while those subjected to mannitol or salt appeared only slightly paler than controls (**Figure 7A**). Quantification using GI confirmed that mannitol and sodium chloride treatments caused moderate but significant reduction in cotyledons greenness, whereas hydrogen peroxide induced a more severe decline (**Figure 7B**). Respiration rates remained statistically unchanged across all treatments (**Figure 7C**). However, the photosynthetic capacity of the stressed plants was markedly reduced compared to those under control conditions. Notably, seedlings exposed to hydrogen peroxide-treated exhibited no detectable photosynthetic oxygen production. Salinity and osmotic stress reduced photosynthetic capacity by around 40 % and 25 %, respectively, compared to control plants (**Figure 7D**).

Finally, we found a strong positive correlation between photosynthesis and the GI values under stress conditions (**Figure 7E**), suggesting that the decline of photosynthetic capacity is, at least in part, associated with stress-induced chlorosis. Together, these results confirm that this oxygen evolution-based approach can sensitively detect stress-induced reductions in photosynthetic function in arabidopsis seedlings.

**Figure 7.**
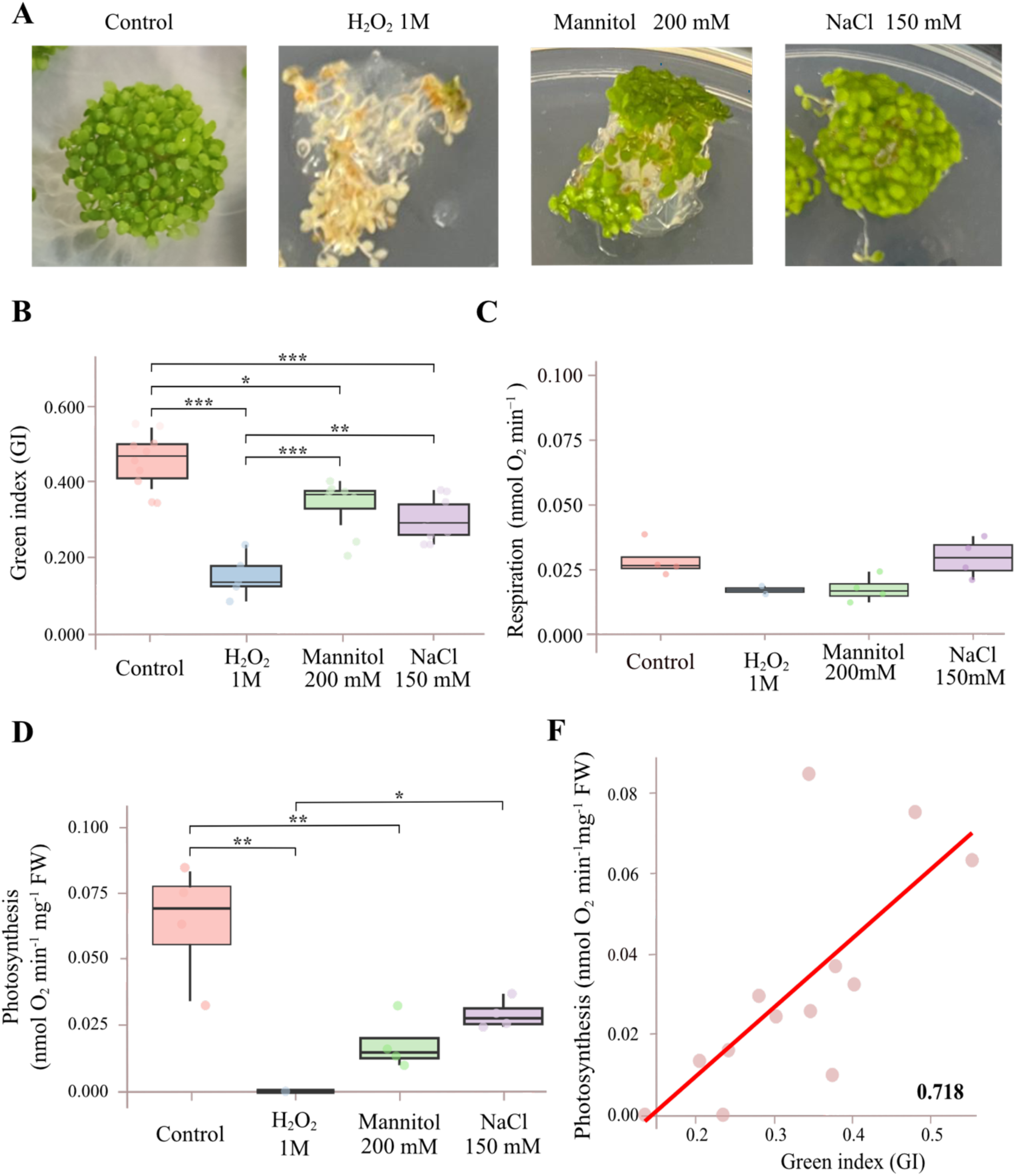
Impact of abiotic stress (osmotic, salt and oxidative) on respiration and photosynthesis. **A.** Phenotypes of one week-old plants grown under MS media, or the last two days on 1M H2O2, 200 mM mannitol or 150 mM NaCl. **B.** Green index values of plants in A. Asterisks represent significant differences according to ANOVA test (* p < 0.05, ** p < 0.01, and *** p < 0.001, n ≈ 10). **C.** Respiration and **D.** Photosynthesis determined by oxygen evolution, of seedlings under control and different abiotic stress treatments (1M H2O2, 200 mM mannitol or 150 mM NaCl). Boxes represent the mean, horizontal lines the median and vertical lines the standard deviation. Dots represent the independent replicates. Asterisks represent significant differences according to ANOVA test (* p < 0.05, ** p < 0.01, and *** p < 0.001, n ≈ 4). **E.** Correlation between GI and photosynthesis. Pearson correlation coefficient is presented in bold in the right bottom corner.

## 4. DISCUSSION

### 4.1. Oxygen electrodes for measuring respiration and photosynthesis

Plant respiration and photosynthesis are central metabolic processes in plant growth and development, both involving in the exchange of O_2_ and CO_2_. Most methods for quantifying these processes are based on detecting changes in gas concentration (Hunt., 2003, Calzadilla et al., 2022) and typically require large plant samples. Here, we evaluated a liquid-phase oxygen electrode system (Oxytherm+P, Hansatech) for real-time measurement of oxygen evolution in arabidopsis seedlings, enabling the estimation of respiration and photosynthesis within minutes. Although oxygen evolution measurements have long been used to assess photosynthesis in system such as arabidopsis leaf discs (Walker and Osmond, 1986; Popova et al., 2018), protoplasts (Hampp et al., 1986), arabidopsis isolated chloroplasts or thylakoids (Delieu and Walker., 1972, Walker and Osmond, 1986; Kansy et al., 2017), and algal culture (Burgess and Davies., 2024), their application to intact plants remains limited. Building on our previous use of this approach in de-etiolated arabidopsis seedlings (Wijerathna-Yapa et al. 2021), we explored the capacity of the Oxytherm+P system (designed specifically for photosynthetic studies) to measure oxygen evolution in small whole-plant samples.

A key advantage of this method is the linear relationship between fresh weight and oxygen evolution (**Figure 1B and C**), allowing flexibility in sample size without compromising data comparability. This enables normalization per gram of fresh weight and eliminates the need for sample mass standardization. To confirm that the observed oxygen fluxes reflect true respiration and photosynthesis, we employed pathway-specific inhibitors. Treatment with potassium cyanide (KCN), which inhibits cytochrome c oxidase, reduced dark-phase oxygen consumption by around 60 % (**Figure 2A**), indicating that most oxygen consumption is due to cytochrome oxidase-dependent respiration. The remaining oxygen consumption is likely attributed to alternative oxidase (AOX), which can bypass complex II and IV to reduce oxygen directly to water (Mclntosh, 1994). Specific AOX genes are known to be induced at the transcriptomic and proteomic level by complex III or complex IV failure (Vanlerberghe et al., 1995 and Karpova et al., 2002), suggesting that AOX could be responsible for most of the oxygen consumption we observed even after the inhibition of cytochrome c oxidase by KCN. To validate the photosynthetic component of the light-phase oxygen evolution, we applied paraquat, an herbicide that blocks photosynthesis by accepting electrons from photosystem I blocking NADPH formation (Qian et al., 2009). Paraquat treatment led to a marked (∼75 %) reduction in oxygen production (**Figure 2B**), confirming that the measured light/dependent oxygen production reflects photosynthetic activity.

Most photosynthetic organism’s own mechanisms for the active uptake of inorganic carbon used by the ribulose-1,5-bisphosphate carboxylase/oxygenase (Rubisco) to metabolize into organic carbon components during the carbon fixation cycle of the photosynthesis process. The addition of sodium bicarbonate (NaHCO_3_) has shown a positive effect on the photosynthesis efficiency on photosynthetic organisms (Salbitani et al., 2020). Here, we used NaHCO_3_ on the measuring buffer as an extra source of inorganic carbon to support plant photosynthesis. The addition of freshly prepared 1 mM of NaHCO_3_ while measuring oxygen evolution at light conditions enhanced over 6-folds the photosynthesis measured in the Clark electrode (**Figure 3**). These results align with previous findings in *Chlamydomonas*, where photosynthetic rates follow a Michaelis–Menten kinetics with bicarbonate concentration (Burgess and Davies., 2024). This has been another strategy to confirm that the oxygen production observed in the Clark electrode is associated with photosynthesis.

Light quantity (quantum flux density) and quality (spectral composition) availability are key determinants of autotrophic growth (Coe and Lin., 2024). There is a well-defined relationship between the amount of irradiance absorbed by plants, the electron transport rate, and Rubisco activity (von Caemmerer., 2000). We recorded a linear increase in oxygen evolution with light intensity up to ∼600 µmol photons m^-2^ s^-1^, followed by a plateau at higher intensities (**Figure 4**), in agreement with established photosynthetic light response curves (Sun and Wang., 2018). At low light intensities, oxygen production is assumed to increase linearly with light, whereas at high intensities a plateau is reached, and photosynthesis becomes independent of light as CO_2_ assimilation is limited by Rubisco activity (Gauthier et al., 2018). Exposing plants to high light intensities above those experienced in the normal growth environment could lead to photoinhibition or photodamage that lead to inhibit photosynthesis (Powles and Thorne., 1981, Ragni et al., 2008). Although our seedlings were grown at 100 μmol m^-2^ s^-1^, they remained metabolically active under brief exposure to much higher light intensity (up to 4500 μmol m^-2^ s^-1^), with no clear evidence of photoinhibition.

Together, these results demonstrate that oxygen evolution measurements using the Oxytherm+P system provide a trustable and physiologically relevant readout of photosynthetic and respiratory activity in arabidopsis seedlings. The method is sensitive to electron transport inhibitors, carbon availability, and light intensity.

### 4.2. Using oxygen electrodes measurements to determine respiration and photosynthesis under different developmental and stress conditions

Light is one of the major environmental factors regulating plant germination, growth, and development. After germinating in soil, seedlings grow heterotrophically in the dark via a process referred to as skotomorphogenesis or etiolation. Etiolated seedlings lack chloroplasts and instead they contain etioplasts (non-green plastids) that do not contain chlorophyll or stacked thylakoid membranes, and hence no photosynthesis takes place (Armarego-Marriott et al., 2020). Etiolation can be prolonged over time by the absence of light after germination and even during the growth of seedlings, where plants develop remarkable morphological, physiological, and biochemical changes (Jedynak et al., 2022). Upon illumination, etiolated seedlings experience a transition from heterotrophic to photoautotrophic growth, since their genetic program switches inducing chlorophyll biosynthesis and, thus, the photosynthetic activity. The manipulation of the light intensity, duration, or the spectral composition of growth conditions have been reported to impact in the photosynthesis activity of different higher plants (Johkan et al., 2012, Wijerathna-Yapa et al., 2021, Li et al., 2022). Here, using our methodology to quantify photosynthesis, we were able to detect differences when plants experience different grades of photomorphogenesis (**Figure 6**). Therefore, this methodology can be used to evaluate the photosynthesis status of diverse mutants related to the photomorphogenesis process or the growth under different treatments associated with light quality, quantity, or photoperiod.

Diverse types of abiotic stresses confer serious damage on the photosynthetic machinery of photosynthetic organisms directly or indirectly where the reduction of net photosynthetic rate strongly decreases (Signorelli et al. 2013; Calzadilla et al. 2016). Alterations such as inhibition of electron or proton transport chain, degradation of chloroplasts thylakoid membrane, and inhibition of chlorophyll biosynthesis have been described (Sharma et al., 2019, Muhammad et al., 2021).

Salinity substantially reduces the overall photosynthetic capacity (Hnilickova et al., 2021), specifically, salinity-induced osmotic stress reduces photosynthesis via the ionic effect on the structure of subcellular organelles and the inhibition of metabolic processes. Additionally, oxidative stress causes reactive oxygen species production causing damage at the photosynthetic machinery (Foyer and Shigeoka., 2011). Regarding the impact of the abiotic stress in the cellular respiration, we did not find changes on this metabolic pathway using our methodology (**Figure 7C**). The impact of abiotic stresses on plant cellular respiration depends on the type, duration, and magnitude of the type of abiotic stress (Flexas et al., 2005, Dey et al., 2021). Most likely, the time we have induced the oxidative, osmotic, and salt stress was not sufficient to observe changes. Moreover, we have detected a strong reduction of photosynthesis rates when arabidopsis seedlings experience two days of salt, osmotic, or oxidative stress (**Figure 7D**). This methodology presents the substantial lead to measure photosynthetic rates as early as a week post-germination and it is relatively sensitive to detect photosynthetic changes when stress is induced in short periods of times. Moreover, we have detected a positive correlation between the greening of seedlings leaves with the photosynthesis rates when plants were exposed to different abiotic stress. The greening of leaves, determined as GI, has been correlated with chlorophyll content (Signorelli et al., 2023), and it is well known that chlorophyll content is directly linked to the photosynthetic activity due to their crucial role in the photosynthetic-light dependent reactions (Sestak., 1963). We noticed that the GI and photosynthesis positive correlation of plants under different light treatments (**Figure 6E**) was smaller to the one of plants under diverse abiotic stress (**Figure 7D**). This is likely attributed to the fact that in the de-etiolated treatment, the greening occurs while seedlings are not yet photosynthetically active, therefore a biphasic response is observed. In the first phase, while greening increases the changes in photosynthesis are minimal, whereas after 24 h of light exposure photosynthesis increases more rapidly resulting in a stronger correlation with the GI parameter (**Figure 7F**).

Temperature strongly influences photosynthetic performance, with both the optimum temperature for photosynthesis and the capacity for acclimation varying among plant species and even ecotypes. In arabidopsis, the optimum temperature for photosynthesis has been reported to be around 22 °C (Thingnaes et al., 2003). At supra-optimal temperatures, various components of the photosynthetic machinery, including membrane fluidity, pigment stability, and enzyme activities, are affected. For example, the oxygen-evolving complex of photosystem II can become inactivated at temperatures exceeding 42 °C (Yamane et al., 1998). Additionally, in C3 plants, photosynthetic inhibition at elevated temperatures is often attributed to reduced carbon assimilation, particularly through decreases in Rubisco carboxylation efficiency and RuBP regeneration (Slattery and Ort, 2019). In our study, we evaluated respiration and photosynthesis at different temperatures during oxygen evolution measurements in the electrode chamber (Figure 7). Although we observed changes in both processes across the temperature range tested, these differences were relatively moderate. It is likely that stronger effects could be detected if seedlings were pre-acclimated to different growth temperatures prior to measurement. Temperature acclimation is known to influence mitochondrial respiration and photosynthetic capacity, with acclimated plants showing greater adjustment of metabolic rates to their growth conditions (Hoffschröer et al., 2024). Future work incorporating such acclimation steps could provide deeper insights into the temperature sensitivity of oxygen evolution in *A. thaliana* seedlings.

### 4.3. Advantages, disadvantages, and considerations for measuring respiration and photosynthesis using an oxygen electrode

This method provides valuable insights into different plant science fields where the metabolic processes of photosynthesis and cellular respiration need to be determined. For example, measuring respiration can be informative of the growth and metabolic activity of non-photosynthetic organs such as roots (Sharma et al. 2011) or axillary buds (Velappan et al. 2022). The liquid-phase oxygen electrode system by the Clark electrode represents an effective, relatively low-cost alternative, and is simple to use in terms of quantifying photosynthesis and cellular respiration of living plants. There are several key advantages associated with the application of this method, that we would like to highlight. i) samples do not need to weigh the same. Although we recommend using the samples within the same range of weight (e.g. 20-50 mg or 80-120 mg) to avoid introducing other variables in the experiment if the researchers do not have the same amount of seedlings for distinct genotypes, different weights can be used as photosynthesis is expressed by fresh weight. ii) The Clark electrode cost is relatively low and access to the instrument is easy since several commercial companies offer it, if we compare with the use of other methodologies such as infrared gas analyser or DUAL-PAM. iii) This protocol is notably fast, providing within minutes rapid measurements of either respiration or photosynthesis in living seedlings. iv) Our methodology using a Clark electrode represents the first approach to measure photosynthesis in living seedlings, making it highly valuable for subsequent analyses and future measurements on the plant material.

Additionally, certain disadvantages might limit the applicability of this methodology. For example, i) we performed all the measurements in a volume of about 2.2 mL (2 mL of measuring buffer, plus the volume of the seedlings) meaning that samples are limited to small volumes. This is a limitation to use this methodology within living plants in bigger size than *A. thaliana* such as crops or legumes. An alternative to sort these limitations is the extraction of protoplast or thylakoid membranes or the generation of leaf discs. ii) The electrode sensor is sensitive to contact with plant material, the samples must not touch the electrode surface because noise can be detected. iii) The sensitivity to electric current fluctuations can be challenged in many laboratory facilities. To mitigate this issue, it is advisable to allow the electrode sufficient time to stabilize before starting with measurements or to utilize a current stabilizer. iv) Plants were initially grown *in vitro* and subsequently transferred into the measuring buffer within the vessel of the electrode. Although the plants remained alive throughout our methodology, the transition from controlled *in vitro* conditions to a new and different environment may introduce variables that could influence the results. v) The light intensity is adjustable within the electrode software; however, the light quality cannot be changed. Studies regarding the growth of plants under different or combinational light spectrum cannot be performed using this method.

Finally, some considerations represent some factors or precautions to ensure the method is applied effectively and in the right circumstances. For example, i) the electrode should include control of temperature and light intensity. ii) while we observed that photosynthetic activity linearly increases with the fresh weight (**Figure 1B and C**), this weight must be between 20 mg to 200 mg. Lower weights mean that the number of seedlings is not enough to generate a floating seedling clump that remains at the upper phase of the buffer during the measurements. When only a few loose seedlings are present they tended to sink and touch the electrode causing noise in the measurements. On the other hand, when the number of seedlings is above 200 mg, the linear relationship between fresh weight and photosynthesis is lost, probably due to too many seedlings shielding some seedlings from the light. iii) we showed that CO_2_ concentrations in the solution dramatically affected the photosynthetic activity (**Figure 3**). Given that the CO_2_ levels dissolved in the solution depend on the pH, buffering for pH is essential for steady-state photosynthesis measurements. Therefore, we recommend checking the buffer if it is not used for more than a month. Also, sodium bicarbonate must be added fresh every day of measurement as it will tend to release CO_2_ reducing its concentration over time. Also because of the need to add sodium bicarbonate fresh and the strong dependency on photosynthesis with sodium bicarbonate concentration, whenever possible we recommend making all measurements within the same day to avoid comparing data with different buffer solutions. iv) the number of replicates or different treatment conditions to be measured may consume a considerable amount of time using one electrode control unit. The Hansatech Oxytherm+P offers up to 8 individual control units to be linked to a single PC and operated simultaneously from OxyTrace+ software providing a powerful, multi-channel system. v) Be aware that the presence of gas bubbles within the liquid phase effects must be removed. vi) Respiration measurements are slower than photosynthesis and must include a buffer blank for proper calibration.

In summary, we demonstrate that oxygen evolution-based measurements provide a reliable and flexible approach to quantify photosynthesis and respiration in intact arabidopsis seedlings. This method enables metabolic assessments during early development and under environmental stress, offering a useful tool for plant physiological studies when conventional gas exchange techniques are not feasible. While here we only show use in wildtype plants, this technique can be used to assess arabidopsis knockout mutants, CRISPR lines, overexpression lines or different ecotypes alongside genetic studies and other phenotype assessments.

## 5. ACKNOWLEDGMENTS

We thank Dr. Mohammad Mukarram for the rich discussion and general comments about our manuscript, and our funding agencies. This project was funded by a grant to SS by the CSIC I+D groups program (Uruguay), group name: “Food and Plant Biology” and group number “883431”. SS and FS are members of the SNI (National Research System, Uruguay) and researchers of PEDECIBA, Uruguay. AB owns a scholarship from Comisión Académica de Posgrado, Universidad de la República, Uruguay. AHM is supported by the Australian Research Council (FL200100057; CE230100015).

## 6. AUTHOR CONTRIBUTION

SS, AHM, and FS conceived the original idea, scheme, and planned all experiments. FS and CC executed the experimental assays. FS and SS analyzed the data, and together with AB and CC created the plots and figures. SS contributed to funding acquisition. All authors contributed to reviewing the literature, wrote, and approved the submitted version.

